# Emergent primordial heritability in short random RNA pools

**DOI:** 10.64898/2026.06.14.732136

**Authors:** Jiro Kakizaki, Alika Andjani Widada, Norikazu Ichihashi, Ryo Mizuuchi

## Abstract

The emergence of heritable molecular information is a fundamental step in the origins of life. Prior to the evolution of defined self-reproducing RNAs in the RNA world hypothesis, mixtures of primordial oligonucleotides may have supported more rudimentary forms of information propagation at the population level. To explore this possibility, we investigated the long-term dynamics of fully random short RNA pools under serial transfer conditions. Across multiple populations undergoing iterative nonenzymatic recombination and ligation, overall sequence compositions changed dynamically; however, sequence and structural traits emerged and exhibited modest yet detectable persistence, indicative of a primordial form of heritability. These traits were largely associated with terminal RNA regions, which tend to be unstructured and accessible for subsequent templating reactions, consistent with the enrichment of complementary sequences. The role of complementarity in shaping sequence composition was further confirmed using designed RNAs. Together, these results suggest that fully random RNA mixtures can exhibit a measurable, albeit weak, capacity to propagate newly generated information in the absence of enzymes, potentially representing an early step toward the emergence of genetic systems.

## Introduction

On primordial Earth, a transition must have occurred from random chemical reactions to organized, self-propagating systems. Such systems can be characterized by their heritable traits, that is, the capability for information transmission that enables reproduction and adaptive evolution. In living organisms, these traits are rooted in the replication of genetic molecules. Similarly, the earliest self-propagating entities may have been based on RNA, as proposed by the RNA world hypothesis (1, 2). Because RNA can function both as an information carrier and as a catalyst, an RNA molecule capable of catalyzing its own reproduction could have emerged, setting the stage for the subsequent evolution of complex biological systems. Inspired by this idea, a growing body of work has attempted to engineer RNA sequences that undergo complete or partial self-reproduction, either through polymerization (3–5) or through more primitive ligation or recombination of RNA fragments (6–8). However, these RNAs are typically complex and incompatible with prebiotically accessible short (∼20-nucleotide (nt)) random oligonucleotides (9, 10). Moreover, even if a short RNA could reproduce (8), its persistence would be challenged within prebiotic, highly diverse oligonucleotide pools that dilute specific substrate RNAs and consume functional sequences through numerous side reactions (11). Thus, a more rudimentary mode of RNA propagation that can spontaneously arise within pools of short random oligonucleotides may have preceded the evolution of efficient self-reproduction by a defined RNA sequence.

A number of studies have proposed that mixtures of diverse, interacting RNAs, each facilitating the synthesis of specific sequences, could collectively propagate via autocatalytic network formation (12–15). In such scenarios, emergent traits at the population level would be maintained over iterative reaction cycles while overcoming dilution or decay, thereby representing a form of primordial heritability (16). A computer simulation based on random ligation among RNA oligonucleotides supported this idea by demonstrating the inheritance of sequence patterns over time (17). Empirically, the emergence of primordial heritability has been explored primarily in non-RNA chemical systems, such as amino acid mixtures and lipid vesicles (18–20). For example, cycles of random amino acid condensation selected specific compositional profiles that persisted against dilution for short periods (18). However, it remains unclear whether a primordial RNA mixture alone could exhibit such heritability, potentially contributing to the establishment of early genetic systems. A related possibility has been examined only in a system employing a polymerase ribozyme, in which emergent sequence biases propagated within random RNA oligonucleotide pools (4).

In this context, RNA oligonucleotides can explore sequence space nonenzymatically via template-directed polymerization, ligation, and recombination (10, 21–27). Several studies have examined these processes in short random RNA pools over short timescales (8, 28–30). For example, we and others demonstrated the spontaneous assembly of 20-nt random RNA oligonucleotides through intermolecular ligation and recombination, resulting in reorganized sequence populations with slight nucleotide biases (8, 30). In these systems, both ligation and recombination likely relied on a 2′,3′-cyclic phosphate (>p), a prebiotically plausible form of RNA activation (31–33); ligation occurs following pre-activation of RNA with >p, whereas recombination proceeds via RNA hydrolysis that generates >p, followed by ligation (21, 23). Our previous study showed that such reactions in random RNA pools can be detected rapidly, within a few days (8), opening the possibility of investigating long-term RNA oligonucleotide dynamics, including the emergence of heritable information.

Here, we examined the long-term dynamics of short random RNA pools under serial transfer conditions. Across multiple RNA populations undergoing iterative nonenzymatic ligation and recombination, overall sequence populations changed dynamically over time, yet sequence and structural traits emerged and persisted for extended periods, suggesting a primitive form of heritability. The strongest contributions to heritable traits were observed in the terminal regions of RNA sequences, which are often located outside intramolecular structures and thus remain accessible to other RNAs for template-directed reactions. Consistently, sequences complementary to these regions were repeatedly enriched, and experiments with defined RNA molecules confirmed the role of complementarity in directing the compositions of random RNA pools. Taken together, fully random primordial RNA mixtures exhibited a measurable, albeit weak, capacity to propagate newly generated information in the absence of enzymes.

## Materials and methods

### RNA preparation

N_20_, N_14_, Semi_AG, and Semi_CU, as well as N_20_ and N_14_ bearing a 3′-terminal monophosphate (N_20_-p and N_14_-p), were purchased from Integrated DNA Technologies (IDT). N_20_ and N_14_ containing a 2′,3′-cyclic phosphate (N_20_>p and N_14_>p) were prepared as described previously (8). Briefly, N_20_-p and N_14_-p were treated with 20 mM 1-(3-dimethylaminopropyl)-3-ethylcarbodiimide hydrochloride (EDC·HCl; Nacalai Tesque) in 300 mM MES buffer (pH 5.5) at 37 °C for 30 min. The resulting N_20_>p and N_14_>p were purified by ethanol precipitation.

### Incubation of random RNA

RNA samples (50 µM N_20_, N_20_>p, N_14_, or N_14_>p) were incubated in 100 mM MgCl_2_ and 50 mM Tris-HCl (pH 8.0) at 22 °C for 2 days. In a subset of experiments, N_14_>p was co-incubated with 0.08–0.8 µM Semi_AG or 0.2–2 µM Semi_CU. Prior to incubation, all RNAs were heated at 65 °C for 2 min and cooled to room temperature. After incubation, an aliquot of the reaction mixture was mixed with four volumes of quenching buffer (50 mM EDTA, pH 8.0, 90% formamide, 0.025% bromophenol blue). Samples were then heated at 95 °C for 2 min, cooled on ice for 1 min, and analyzed by denaturing polyacrylamide gel electrophoresis (PAGE) on a 20% polyacrylamide / 8 M urea gel in 1×TBE buffer. Gels were stained with SYBR Gold (Thermo Fisher Scientific), and RNA bands were visualized using FUSION-SL4 (Vilber-Lourmat). The obtained images were analyzed using ImageJ (NIH).

### Serial transfer experiment

After incubating a random RNA pool for 2 days as described above, an aliquot was diluted 5-fold in the same buffer (100 mM MgCl_2_, 50 mM Tris-HCl, pH 8.0) containing 50 µM of the original random RNA pool. This procedure was repeated for approximately 30 rounds. Samples were quenched at every 1–3 rounds as described above and stored at −80 °C until use. To remove terminal phosphates from N_20_>p- and N_14_>p-derived RNAs, ∼100 pmol of RNA was ethanol-precipitated and treated with 10 U T4 polynucleotide kinase (New England Biolabs) at 37 °C for 1 h. RNA samples were then analyzed by 20% denaturing polyacrylamide gel electrophoresis and visualized after SYBR Gold staining, as described above. RNA products corresponding to approximately 15–45 nt (N_14_ and N_14_>p-derived products) or 21–45 nt (N_20_ and N_20_>p-derived products) were excised from the gel and ethanol-precipitated.

For high-throughput sequencing (HTS) analysis, the collected RNA products were converted to cDNA libraries using the SMARTer smRNA-Seq Kit for Illumina (Takara), as described previously (8). In this procedure, RNA products were 3′ polyadenylated, reverse-transcribed, and PCR-amplified. For multiplexed sequencing, indexed primers from the SMARTer RNA Unique Dual Index Kit (Takara) were used during PCR amplification. The amplified libraries were electrophoresed on a 15% native polyacrylamide gel (SuperSep DNA, Fujifilm) in 1×TBE buffer, visualized by SYBR Gold staining, excised, and purified using NucleoSpin Gel and PCR Clean-up (Takara). Libraries derived from a single transfer experiment were pooled and sequenced on a DNBSEQ-G400 platform (PE100) at BGI Hong Kong.

### Sequence processing

Raw demultiplexed sequence reads were trimmed using cutadapt (34) according to the manufacturer’s protocol for the SMARTer smRNA-Seq Kit for Illumina (Takara). The remaining reads were then quality filtered as shown in Supplementary Fig. S1. First, cutadapt was used to trim low-quality bases using a Q30 threshold, and only forward reads showing perfect complementarity to reverse reads were retained. Next, BLAST+ v2.16.0 (35) was used to identify sequences sharing homology with a custom database consisting of primer sequences, electrophoresis markers, and reads obtained from control libraries prepared using the same experimental protocol without input RNA. The search was performed with default parameters with the addition of “-strand plus” to exclude reverse-complement matches, and sequences showing > 90% identity and query coverage were removed. To further exclude potential contaminants, enriched sequences whose frequency ranked within the top 200 and exceeded 5×10^-7^ at each round were searched against the NCBI core nucleotide database (core_nt; downloaded between September 15 and 19, 2025) (36) (Supplementary Fig. S2). The homologous sequences identified in this search were used to construct a potential contamination reference database, and sequences showing >90% identity and query coverage to this database were further removed from the experimental dataset. From the remaining N_14_-, N_14_>p-, N_20_-, and N_20_>p-derived sequences, 100,000 sequences were randomly selected within length ranges of 18–30 nt, 18–30 nt, 25–45 nt, and 25–45 nt, respectively, and subjected to detailed analyses.

### Sequence composition analysis

Population diversity was quantified using normalized Shannon population entropy, calculated as described previously (37):

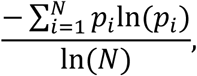

where *p_i_* represents the frequency of *i*-th sequence and *N* is the total number of unique sequences in the population.

To assess heritable traits, the frequencies of all *k*-nucleotide (*k*-mer) patterns were calculated for each of *N* sequences, generating an *N* × 4*^k^*data matrix. Each column was standardized and subjected to principal component analysis (PCA) using the Python package scikit-learn. For each population, principal component scores were calculated, and the first 3 × 4*^k^*^-1^ principal components were retained, generating an *N* × (3 × 4*^k^*^-1^) dataset. For *k* = 1–4, the cumulative explained variance approached 1 (Supplementary Fig. S3), and the resulting principal component scores for *k* = 1–3 approximately followed a multivariate normal distribution (Supplementary Fig. S4). Accordingly, for each round *i*, the distribution of 3-mers, *P_i_*(***x***), was approximated by a multivariate normal distribution with mean vector μ*_i_* and covariance matrix Σ*_i_* estimated from the principal component score vector ***x***. The dissimilarity between distributions from rounds *i* and *j* was quantified using the Jensen–Shannon (JS) divergence (38), *D_JS_*:

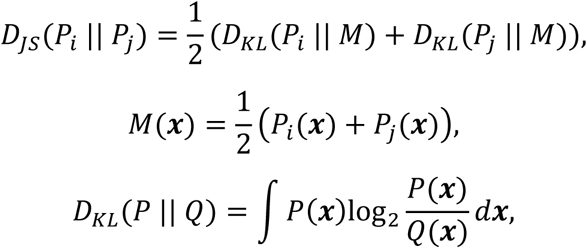

where *D_KL_* denotes the Kullback–Leibler divergence. *D_JS_* takes values between 0 and 1, with smaller values indicating greater similarity between the two distributions.

To assess the statistical significance of the observed apparent heritability, pairwise *D_JS_* values were calculated between all rounds to construct a divergence matrix *d_JS_* for rounds *i* and *j*. The mean divergence between consecutive rounds (*d^C^*) and that between non-consecutive rounds (*d^NC^*) were then calculated, and their difference was defined as the statistic *e*:

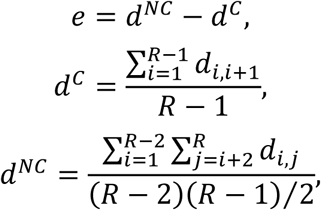

where *R* denotes the total number of rounds. The null hypothesis that the observed *e* could arise by chance regardless of the ordering of rounds was tested by randomly permuting the round labels to generate new divergence matrices, followed by recalculation of *e* (*e_per_*) for each permutation. After *X* (= 100,000) permutations, the null distribution of *e* was obtained, and a right-tailed test was performed. The *p*-value was calculated as *p* = (*S* + 1)/(*X* + 1), where *s* denotes the number of permutations in which the recalculated statistic exceeded the observed value *e_obs_*.

To assess the contribution of individual 3-mers, the columns corresponding to each 3-mer in the original *N* × 4^3^ 3-mer frequency matrix were randomly shuffled, and the divergence matrix was recalculated. To evaluate positional contributions, 3 nucleotides at a given position from either the 5′ or 3′ terminus were removed from each sequence, and the divergence matrix was recalculated. These procedures generated a new statistic *e_i_*. The contribution of each 3-mer or nucleotide position to heritability was quantified as c = (*e_obs_* − *e_i_*)/*e_obs_*, where larger values of *c* indicate stronger contributions.

### Sequence complementarity analysis

To assess sequence complementarity, three subsequence populations were prepared: original product sequences, 6-nt unpaired terminal regions defined based on predicted minimum free energy (MFE) structures as described below, and all 6-nt terminal regions. Unpaired termini shorter than 6 nt were excluded. For each sequence, the 6-nt regions at both termini were concatenated to generate a 12-nt sequence, as the combined termini may act as a single template. To evaluate sequence complementarity during serial transfer, products from the initial rounds were excluded from the analysis. The number of sequence pairs containing at least one fully complementary 6-nt region was then counted by comparing original products with each of the three subsequence populations. The statistical significance of the observed increase in complementary pairs was assessed using a permutation test similar to that described in the previous section. In this analysis, the difference between the mean count for identical or consecutive rounds and that for non-consecutive rounds was defined as the statistic *e*, and the significance of the observed value *e_obs_* was evaluated using a one-sided test with 100,000 permutations.

### RNA structural analysis

RNA secondary structures were predicted as minimum free energy (MFE) structures at 22 °C using the Vienna RNA Package (39). The free energies of the structures (Δ*G*_MFE_) were then obtained. Structural dissimilarities between populations were quantified using the JS divergence between histogram-based ΔG_MFE_ distributions (bin width = 0.5).

## Results

### Long-term incubation of short random RNA pools

As model primordial RNA pools, we prepared fully random 14-nt and 20-nt RNAs (N_14_ and N_20_), as well as their 2′,3′-cyclic phosphate-activated counterparts (N_14_>p and N_20_>p, respectively). Each pool contained approximately 3 × 10^14^ molecules, theoretically covering all possible ∼2.5×10^8^ and ∼10^12^ sequences with redundancy for 14-nt and 20-nt oligonucleotides, respectively. N_14_ and N_20_ were expected to undergo spontaneous recombination, whereas N_14_>p and N_20_>p could undergo both recombination and ligation, generating products of varying lengths. Consistent with our previous study on N_20_ and N_20_>p (8), all four pools generated elongated products, primarily up to approximately twice their original length, after 2 days of incubation at 22 °C in the presence of 100 mM MgCl_2_ (Fig. 1A). Although the increase in N_20_-derived products appeared small on denaturing PAGE, this was likely due to undenatured RNA complexes that formed even without incubation, as they did not amplify in subsequent RT-PCR (8). We next subjected the RNA pools to serial transfer experiments (Fig. 1B). Each round consisted of (i) incubation under the same reaction conditions and (ii) five-fold dilution with the initial random RNA pools to replenish short oligonucleotides. Over approximately 30 rounds, denaturing PAGE revealed no substantial changes in the overall product length distributions (Supplementary Fig. S5). However, these results do not exclude potential changes in sequence composition.

**Figure 1.**
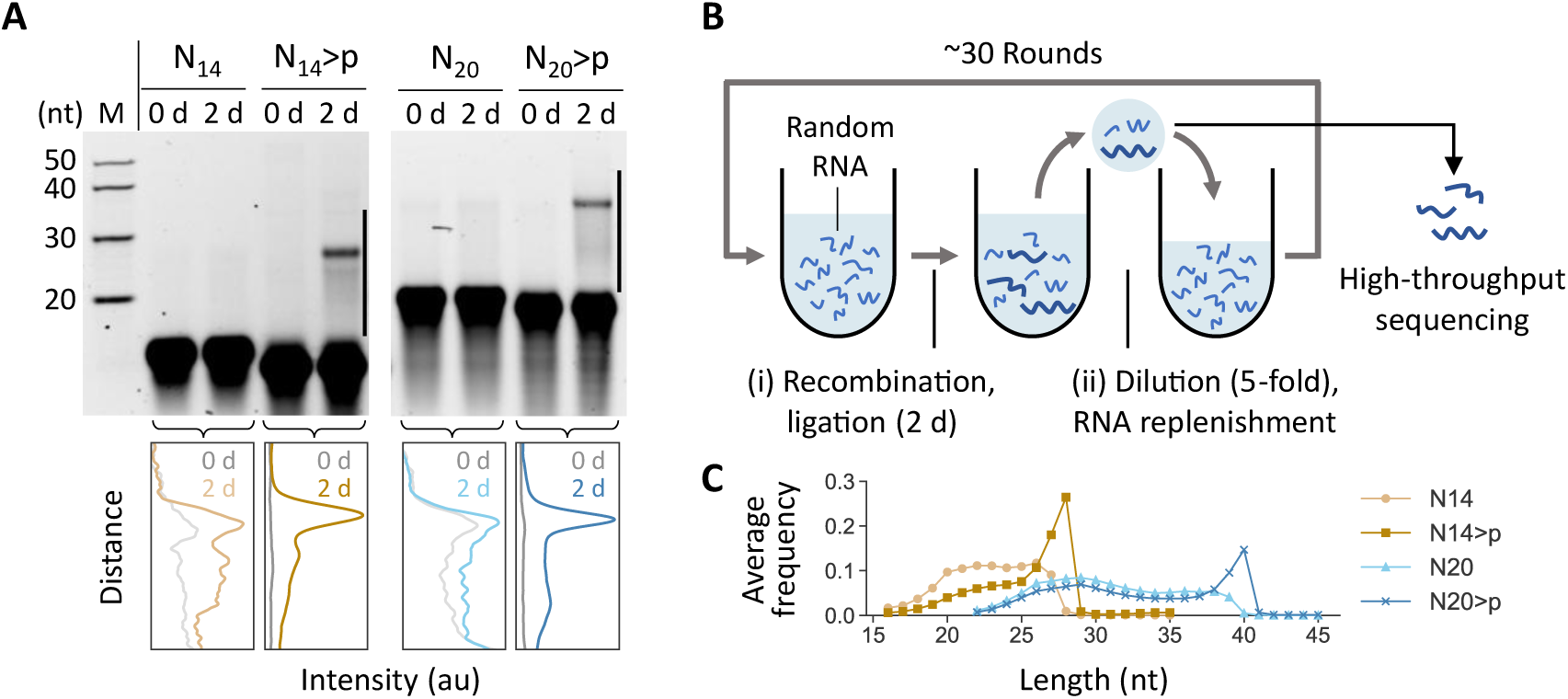
Long-term incubation of short random RNA pools. (A) Incubation of 50 μM N_14_, N_14_>p, N_20_, or N_20_>p for 2 days at 22 °C in 100 mM MgCl_2_ (pH 8.0), analyzed by 20% denaturing PAGE. RNAs with >p migrated slightly faster due to additional negative charge. Band intensities corresponding to the regions indicated by the black lines were quantified for each pool (bottom). Length-dependent differences in staining efficiency were not corrected. (B) Schematic of the serial transfer experiment. (i) A selected RNA pool was incubated under the same conditions for 2 days to induce recombination and ligation. (ii) The population was diluted five-fold while replenishing the initial random RNA pool. Populations after step (i) were subjected to high-throughput sequencing. (C) Product length distributions averaged across all rounds. Products one nucleotide longer than the original lengths (i.e., 15- or 21-mers) were excluded as they may have arisen from the addition of an extra nucleotide to the original RNA pools during template switching.

To examine sequence-level dynamics, elongated products were analyzed by high-throughput sequencing (Fig. 1B). Products ranging from ∼15 nt (N_14_ and N_14_>p pools) or ∼21 nt (N_20_ and N_20_>p pools) to ∼45 nt were excised from denaturing gels every 1–3 rounds. The full-length RNAs were subjected to RT-PCR using SMART (switching mechanism at the 5′ end of the RNA transcript) technology (40), which combines 3′ polyadenylation and template switching during reverse transcription, followed by high-throughput sequencing analysis. Sequence reads were quality-filtered, and potential contaminants were removed (Supplementary Figs. S1 and S2). Recombination between two 14-nt or two 20-nt RNAs was expected to generate products of 15–27 nt and 21–39 nt, respectively, whereas ligation would produce 28-nt or 40-nt products. Consistent with these mechanisms, most reads fell within 18–27 nt (N_14_), 18–28 nt (N_14_>p), 25–39 nt (N_20_), and 25–40 nt (N_20_>p) across all rounds (Fig. 1C and Supplementary Fig. S6). Sharp drops at 28 nt and 40 nt were observed in N_14_- and N_20_-derived products, respectively, whereas corresponding peaks were observed in the N_14_>p- and N_20_>p-derived products. For downstream analyses, we randomly sampled 100,000 sequences at each round from 18–30 nt (N_14_ and N_14_>p) or 25–45 nt (N_20_ and N_20_>p); all analyses were repeated with at least ten independent random samplings. To examine changes in overall sequence diversity, we counted identical sequences observed across different rounds (Supplementary Fig. S7). Identical sequences were rarely detected across rounds in all four pools, indicating that global sequence diversity was largely preserved throughout the transfer experiments. Consistently, Shannon population entropy, a measure of population-level sequence diversity, remained close to 1 throughout the rounds for all four pools (Supplementary Fig. S8).

### Shifts in nucleotide compositions

While individual sequences changed dynamically, distinctive compositional traits may have emerged and persisted through successive transfers. To examine this possibility, we analyzed changes in nucleotide composition of the products across rounds. Compositional shifts with apparent persistence were detected in N_14_-, N_14_>p-, and N_20_>p-derived products (Fig. 2A). For example, in N_14_-derived products, the frequency of G increased by ∼5% at round 2 and remained elevated until approximately round 26. Similarly, in N_20_>p-derived products, the frequency of U increased by ∼7% at round 17 and persisted until the final round. N_14_>p-derived products exhibited oscillatory yet partially consistent compositional dynamics, whereas N_20_-derived products showed clear compositional shifts only during rounds 24–29, which were not consistently maintained.

**Figure 2.**
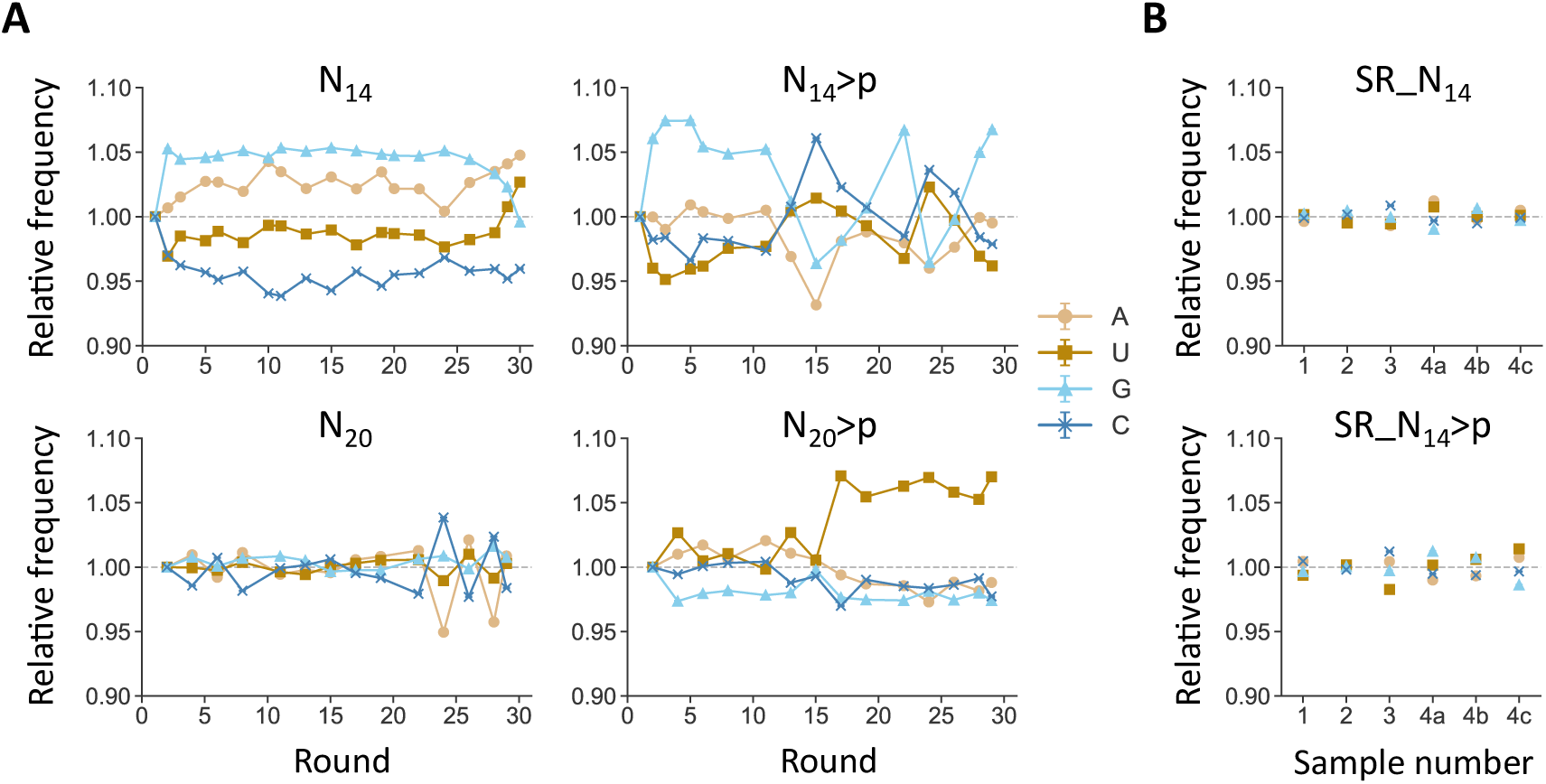
Changes in nucleotide composition during serial transfer. (A) Frequencies of individual nucleotides relative to those in the initial round for N_14_-, N_14_>p-, N_20_-, and N_20_>p-derived products. Error bars (mostly smaller than the markers) indicate standard errors based on random sampling of 100,000 sequences (*n* = 10). (B) Frequencies of individual nucleotides in products derived from single-round incubations of N_14_ and N_14_>p pools (SR_N_14_- and SR_N_14_>p), shown relative to the average across six experiments. Experiments were repeated four times starting from independent incubations of random RNA pools (samples 1–4). For sample 4, library preparation for high-throughput sequencing was performed in triplicate (denoted as 4a, 4b, and 4c).

To assess the significance of the observed compositional shifts, we performed multiple single-round incubations of the N_14_ and N_14_>p populations under the same conditions as the transfer experiments (hereafter denoted SR_N_14_ and SR_N_14_>p, respectively). Across six replicates for each population, changes in nucleotide frequencies relative to the replicate mean were at most 1.2% and 1.7% for SR_N_14_- and SR_N_14_>p-derived products, respectively (Fig. 2B). These values were substantially smaller than the compositional shifts observed during serial transfer, suggesting that nucleotide composition changes in N_14_- and N_14_>p-derived products were associated with the recursive transfer process, involving repeated ligation and recombination coupled with replenishment of random RNA oligonucleotides. Overall, persistent nucleotide compositional changes were observed in multiple populations undergoing serial transfer, consistent with population-level compositional inheritance.

### Shifts in nucleotide patterns

To assess potential heritability of higher-order sequence information, we analyzed the dynamics of nucleotide patterns (subsequences, or *k*-mers). Overall changes in *k*-mer composition were quantified by calculating Jensen–Shannon (JS) divergence (38) between populations at different rounds based on *k*-mer frequency profiles of individual sequences. JS divergence measures dissimilarity between multidimensional probability distributions. Here, it was calculated from *k*-mer frequency vectors (e.g., frequencies of AAA, AAU, etc. for *k* = 3) after principal component analysis (see Materials and methods). We focused on *k* = 3, the largest *k* allowing reliable estimation of JS divergence (Supplementary Fig. S4).

Round-by-round comparison of JS divergence reflected shifts in 3-mer composition (Fig. 3A). Consistent with the nucleotide compositional shifts (Fig. 2), 3-mer features emerged and persisted over multiple rounds. For example, more similar 3-mer compositions were observed between rounds 2 and 26 in N_14_-derived products and between round 17 and the final round in N_20_>p-derived products. To quantify heritable aspects of 3-mer compositions, we compared JS divergence between consecutive rounds and between non-consecutive rounds, as lower divergence is expected between consecutive rounds in heritable populations. Lower divergence was detected between consecutive rounds compared with non-consecutive rounds in N_14_-, N_14_>p-, and N_20_>p-derived products (Fig. 3B). To further assess this apparent heritability, we evaluated the statistical significance of the difference between the mean divergence of consecutive rounds (*d^C^*) and that of non-consecutive rounds (*d^NC^*) by repeatedly shuffling round labels in the divergence matrices, followed by comparison of the observed statistic (*e_obs_*) with values obtained from permuted matrices (*e_per_*) (Fig. 3C). The resulting low p-values for N_14_-, N_14_>p-, and N_20_>p-derived products supported heritable nature of their 3-mer compositions (Fig. 3A). Among the 64 possible 3-mers, nearly all contributed positively to the observed heritability, although those showing stronger contributions differed among pools (Supplementary Fig. S9), suggesting that the heritability reflects broadly distributed sequence features rather than being driven by small subsets of sequences. Divergence between non-consecutive rounds was also greater than that observed in SR_N_14_- and SR_N_14_>p-derived products (Fig. 3B), further supporting the role of recursive transfer in shaping 3-mer compositions.

**Figure 3.**
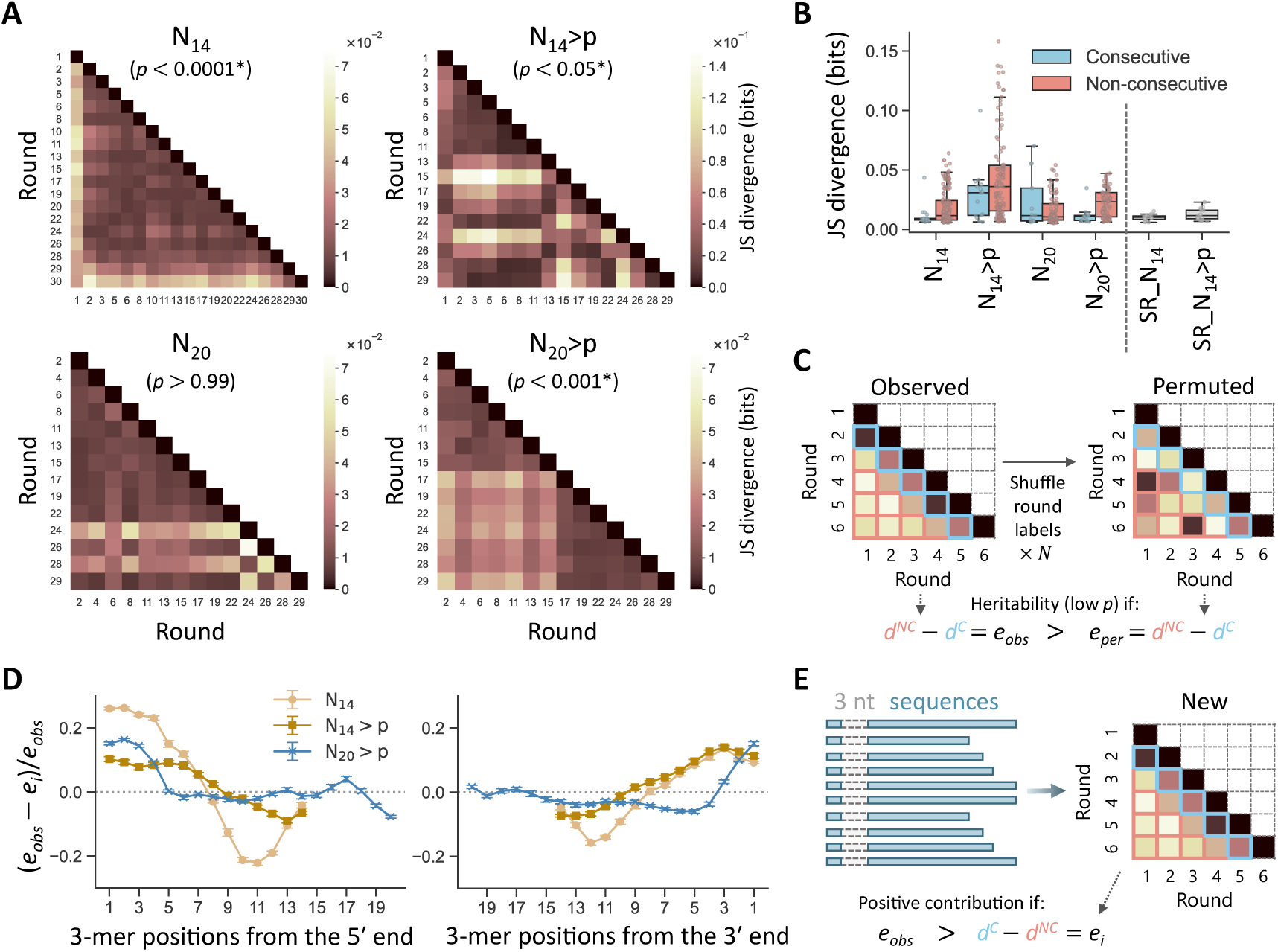
Shifts in 3-mer patterns as heritable traits. (A) Round-by-round comparison of JS divergence based on 3-mer frequency vectors between populations at any two rounds, calculated for N_14_-, N_14_>p-, N_20_-, and N_20_>p-derived products. Average divergence from randomly sampled 100,000 sequences is shown (*n* = 10). (B) Box plots showing distributions of JS divergence between consecutive rounds and between non-consecutive rounds, compared with SR_N_14_- and SR_N_14_>p-derived products. Each data point corresponds to an individual divergence in panel A. (C) Schematic of the statistical evaluation of apparent heritability. p-values calculated from the averaged JS divergence matrices are shown in parentheses in panel A; asterisks indicate *p* < 0.05. (D) Position-dependent contributions of 3-mers to the observed heritable traits in N_14_-, N_14_>p-, and N_20_>p-derived products, quantified as changes in JS divergence after excluding specific 3-mer positions from the 5′ (left) or 3′ end (right). Error bars indicate standard errors from the same random sampling (*n* = 10). (E) Schematic of the quantification of position-dependent 3-mer contributions shown in panel D.

Next, we examined position-dependent 3-mer contributions to the observed heritable traits by quantifying changes in JS divergence after excluding specific 3-mer positions from sequences (Fig. 3D and E). In N_14_-, N_14_>p-, and N_20_>p-derived products, 3-mers near the 5′ and 3′ termini contributed positively, whereas those in internal regions contributed negatively or negligibly. For example, up to the 7th–8th positions from both termini in N_14_- and N_14_>p-derived products exhibited positive contributions. These terminal regions tended to form unpaired dangling ends, whereas internal sites were more likely to be structured (Supplementary Fig. S10). Such unpaired terminal regions may have served as templates for subsequent ligation or recombination, potentially contributing to the observed heritable traits.

To examine the robustness of the emergence of heritable traits, we performed two additional 30-round transfer experiments using newly prepared N_14_ and N_14_>p pools, followed by sequence analysis using the same method (Supplementary Fig. S11). In these experiments, heritable 3-mer dynamics were statistically significant in N_14_>p-derived products but not in N_14_-derived products. Consistent with the initial experiment (Fig. 3D), terminal regions in N_14_>p-derived products exhibited positive contributions to heritability, whereas internal regions showed negative contributions.

### Shifts in structural patterns

To examine changes in traits beyond nucleotide sequences, we analyzed RNA secondary structures. Minimal free energy (MFE) structures were predicted for all products, and their free energies (ΔG_MFE_) were used as a proxy for structural stability. JS divergence was then calculated between ΔG_MFE_ distributions of any two rounds for each pool (Fig. 4A and B). For N_14_-, N_14_>p-, and N_20_-derived products, the resulting JS divergence matrices were similar to those based on nucleotide patterns (Fig. 3A), with correlation coefficients of 0.70, 0.75, and 0.87, respectively. In contrast, N_20_>p-derived products showed weaker correspondence, with a correlation coefficient of 0.39. Notably, N_14_- and N_14_>p-derived products exhibited statistically significant heritable structural dynamics. This observation is consistent with the expectation that sequence composition influences secondary structure. However, N_20_>p-derived products did not show heritable structural dynamics despite detectable 3-mer-based heritability, suggesting that similar structural distributions may arise from distinct nucleotide patterns. N_20_-derived products likewise did not show heritable structural dynamics, consistent with their relatively stable nucleotide patterns.

**Figure 4.**
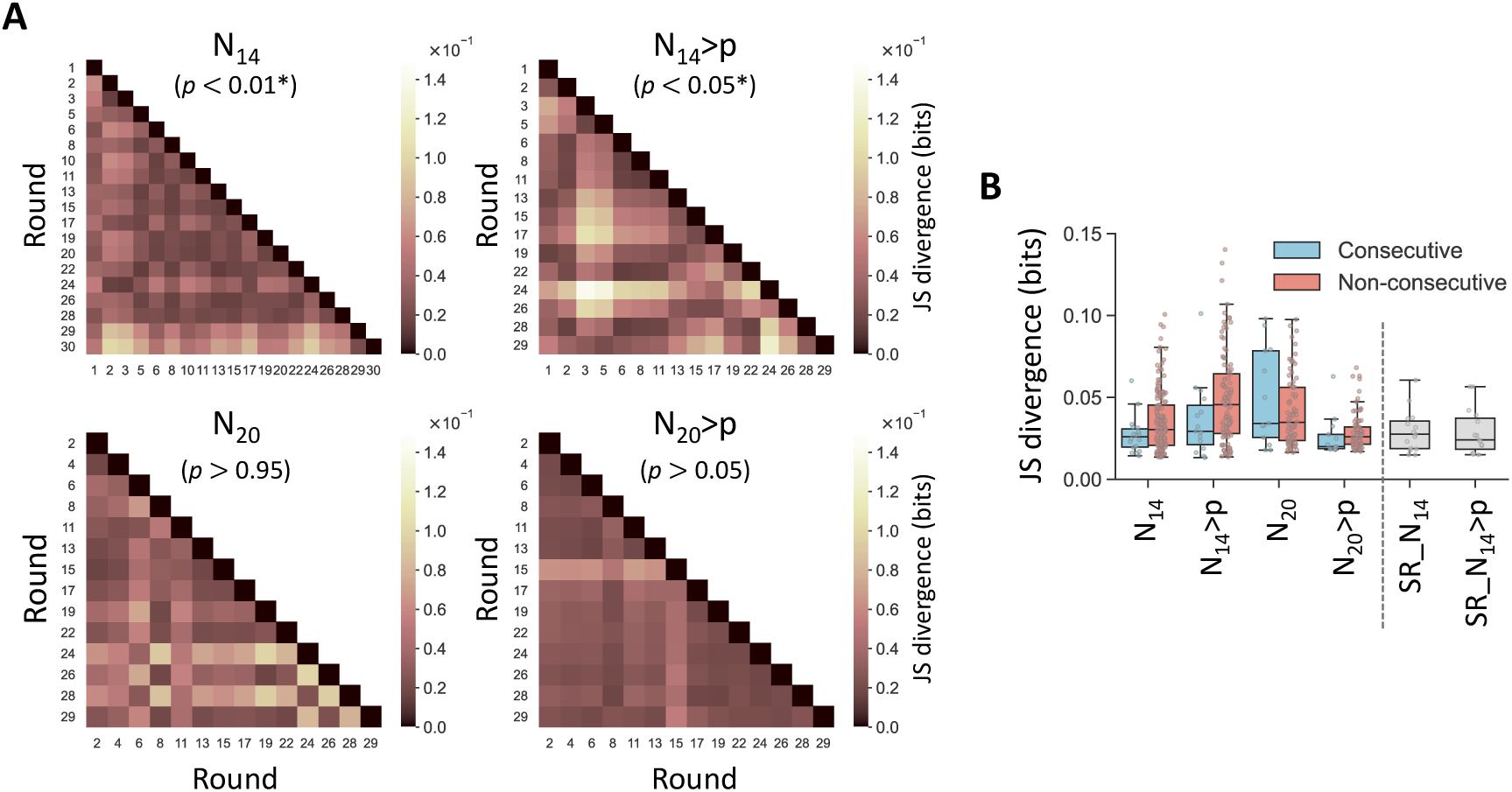
Shifts in structural patterns as heritable traits. (A) Round-by-round comparison of JS divergence based on ΔG_MFE_ distributions between populations at any two rounds, calculated for N_14_-, N_14_>p-, N_20_-, and N_20_>p-derived products. Average divergence from randomly sampled 100,000 sequences is shown (*n* = 10). p-values were obtained as in Fig. 3C and are shown in parentheses; asterisks indicate *p* < 0.05. (B) Box plots showing distributions of JS divergence between consecutive rounds and between non-consecutive rounds, compared with SR_N_14_- and SR_N_14_>p-derived products. Each data point corresponds to an individual divergence in panel A.

### Enrichment of complementary sequences

Motivated by the observation that terminal sequences contributed positively to heritable dynamics and often appeared as unpaired dangling ends, we hypothesized that such accessible regions could serve as templates for subsequent ligation and recombination reactions. Under this scenario, terminal regions of products from identical or consecutive rounds would be expected to display higher inter-sequence complementarity than those separated by larger time gaps. To test this hypothesis, we quantified sequence complementarity by counting complementary 6-nt pairs between subsequence populations across all round combinations, comparing (i) 6-nt unpaired terminal regions (UT) and full-length original product sequences (O) (Fig. 5A). We focused on 6-nt terminal sites because these regions showed relatively high contributions to the observed heritability (Fig. 3D). For comparison, we also analyzed (ii) all 6-nt terminal regions, including both paired and unpaired termini (T), and (iii) full-length original sequences (O).

**Figure 5.**
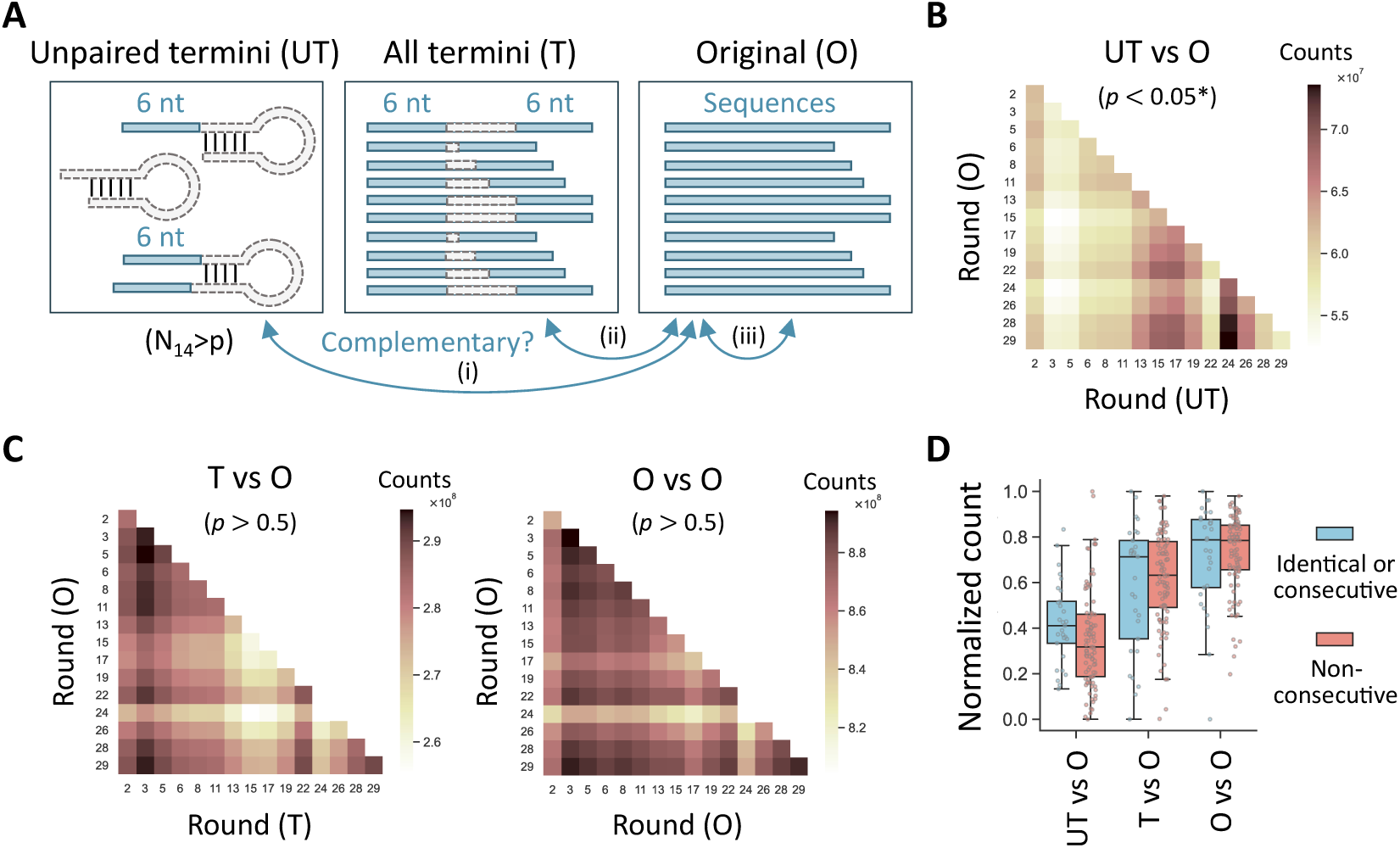
Enrichment of complementary sequences in N_14_>p-derived products. (A) Schematic of sequence complementarity analysis comparing (i) 6-nt unpaired termini (UT), (ii) all 6-nt termini (T), and (iii) full-length original product sequences (O), each against original sequences across all round combinations. (B) Round-by-round comparison of the number of complementary 6-nt pairs between UT and O populations. Average counts from randomly sampled 100,000 sequences are shown (*n* = 10). (C) Same comparison as in panel B between T and O populations (left) or between O and O populations (right). p-values calculated from the averaged matrices are shown in parentheses; asterisks indicate *p* < 0.05. (D) Box plots showing distributions of the number of complementary 6-nt pairs after min–max scaling between identical or consecutive rounds and between non-consecutive rounds.

For N_14_>p-derived products, 6-nt unpaired termini displayed higher complementarity to original sequences between identical or consecutive rounds than between non-consecutive rounds (Fig. 5B and D). The increased complementarity between nearby rounds was statistically significant, as assessed using the same permutation framework applied to the JS divergence analysis (Fig. 3C). In contrast, including paired termini or analyzing full-length sequences abolished this tendency (Fig. 5C and D). N_14_-derived products showed similar tendencies, whereas N_20_>p-derived products did not (Supplementary Fig. S12), possibly because the contribution of terminal regions to heritability was smaller in N_20_>p-derived products than in N_14_>p- and N_14_-derived products (Fig. 3D). These results support a potential role for sequence complementarity in the emergence of heritable traits.

### Complementarity-directed sequence generation

Motivated by the observed potential contributions of unpaired sequence regions to the enrichment of complementary sequences, we further examined whether predefined sequences could selectively generate complementary sequences in random RNA pools. To this end, we performed a single-round transfer experiment using an N_14_>p pool in the presence of semi-random RNAs containing defined sequence regions. We designed semi-random 22-nt RNAs, AG(NNAG)_5_ and CU(NNCU)_5_, which represent potential ligation and recombination products with lower structural propensity than fully random sequences (hereafter denoted semi_AG and semi_CU, respectively) (Supplementary Fig. S13A). After co-incubating 50 μM N_14_>p RNA with 0.8–2 μM of either semi-random RNA, products were analyzed by high-throughput sequencing using the same approach as in the transfer experiments (Fig. 6A). A parallel reaction lacking semi-random RNAs served as a negative control.

**Figure 6.**
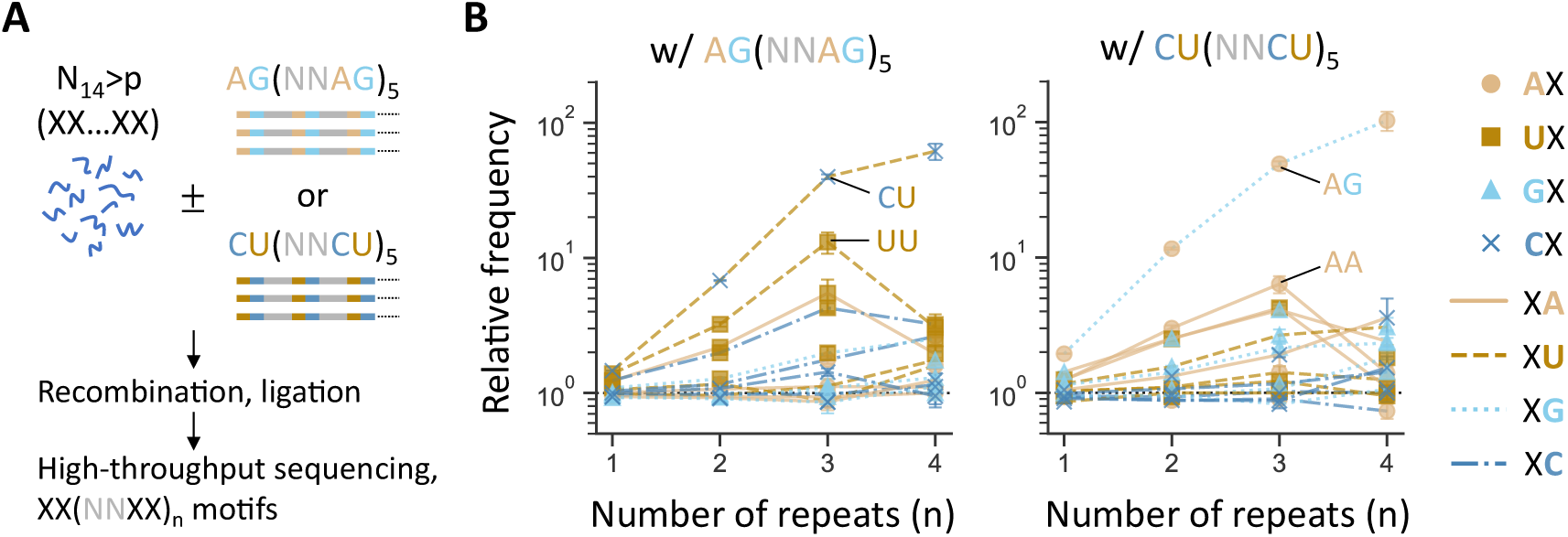
Complementarity-directed sequence generation in N_14_>p pools. (A) Schematic of the experiment combining N_14_>p RNA with semi-random RNAs. (B) Frequencies of XX(NNXX)_n_ motifs detected after incubating the N_14_>p pool in the presence of 0.8 μM AG(NNAG)_5_ (left) or 2 μM CU(NNCU)_5_ (right), relative to those in the absence of semi-random RNAs. Different combinations of marker shapes and line styles represent the 16 possible dimers (XX). Error bars indicate standard errors based on random sampling of 100,000 sequences (*n* = 10).

Among the detected products, we analyzed the distribution of XX(NNXX)_n_ motifs, where XX represents one of the 16 possible dimers (AA, AU, etc.) and n (= 1–4) denotes the number of repeats. In the presence of semi-random RNAs, products containing XX motifs complementary to the fixed AG or CU sequences were selectively enriched, with enrichment increasing with repeat number (Fig. 6B). For example, CU(NNCU)_4_ and AG(NNAG)_4_ were approximately 100-fold more abundant in the presence of semi_AG and semi_CU, respectively, than in the negative control. Even motifs containing only two repeats (CU(NNCU)_2_ and AG(NNAG)_2_) were enriched by approximately 10-fold. XX motifs partially complementary to AG or CU were also well enriched (e.g., UU for semi_AG and AA for semi_CU).

To assess the robustness of this selective enrichment, we repeated the experiments with a 10-fold lower input of the semi-random RNAs (Supplementary Fig. S13B). Sequence analyses revealed similar trends, albeit with slightly reduced enrichment. These results highlight the role of sequence complementarity in directing the composition of RNA populations, representing a primitive form of information propagation.

## Discussion

Our results demonstrate that primordial pools of short random RNAs can exhibit a measurable form of heritability through iterative, prebiotically plausible nonenzymatic ligation and recombination via 2′,3′-cyclic phosphate activation. During serial transfer, newly generated sequences continuously reshaped overall population compositions, as expected from the high diversity of the starting oligonucleotide pools. Despite these dynamic changes, emerging nucleotide patterns and structural features exhibited modest but detectable persistence across generations (Figs. 3 and 4). Such emergent heritability was reproducibly observed in four of six independent RNA pools, indicating that the phenomenon is reasonably robust. In these systems, only a small fraction of sequence information was retained between generations, in contrast to biological replication, which copies complete genetic templates. Nonetheless, this partial retention of molecular information suggests that random RNA populations possess an intrinsic capacity to propagate newly generated features, even if only weakly, potentially representing an early step toward the emergence of RNA-based genetic systems.

Since template-directed ligation and recombination are the primary reactions expected to occur in the random RNA mixtures, these processes likely underlie the observed heritability. Our analysis indicates that heritability largely relies on the terminal regions of reaction products, which tend to be unstructured and thus more likely to remain accessible for subsequent templating events (Figs. 3D and Supplementary Fig. S10). Consistently, nucleotide patterns complementary to these regions were detected relatively frequently in identical or consecutive rounds, and RNA motifs complementary to the input sequences were also enriched (Figs. 5 and 6). Similar enrichment of complementary sequence motifs has been reported in previous studies of protein-catalyzed templated ligation or polymerization of random DNA pools (41, 42). Likewise, simulations of ligation or recombination within virtual random RNA populations have demonstrated the emergence of partial information propagation driven by sequence complementarity (17, 43). Together, these observations support a role for simple RNA–RNA hybridization in information propagation within primordial RNA mixtures.

The random RNA populations subjected to serial transfer exhibited not only primordial heritability but also greater shifts in compositional features of reaction products than untransferred controls (Figs. 2 and 3B). Recursive cycles of product dilution and substrate replenishment can maintain chemical systems out of equilibrium, thereby imposing selection pressures on evolving mixtures. Previous studies have shown that even relatively short recursive cycles can alter product distributions generated by random condensations of chemicals such as amino acids and formamide (18, 44, 45). Similar observations in experiments involving random ligation and recombination of short RNA molecules further underscore the importance of recursive cycling in selecting specific products within largely random mixtures.

While our experimental system is based on RNA chemistry, the emergence of weak yet persistent compositional shifts observed here resonates with theoretical models of early chemical evolution. For example, the graded autocatalysis replication domain (GARD) model (46, 47) describes mutually catalytic lipid assemblies in which compositional inheritance emerges through the growth and division of heterogeneous molecular aggregates, representing a form of self-reproduction. In this framework, environmental perturbations, such as changes in available molecular species, can induce transitions between distinct persistent compositional states, suggesting a primitive form of evolutionary dynamics (48). Similarly, although the random RNA oligonucleotides examined here exhibited only modest shifts and partial persistence of sequence composition, environmental fluctuations could amplify such biases and enable transitions toward more differentiated compositional states. Such dynamics may represent an early step toward the emergence of primitive sequence-based evolutionary dynamics.

## Data Availability

High-throughput sequencing data have been deposited in the NCBI Sequence Read Archive under BioProject accession number PRJNA1474794. A reviewer link is available at xxx. All other data and code relevant to this study are available in the main text, the supplementary materials, or from https://doi.org/10.6084/m9.figshare.32583090.

## Acknowledgements

We thank David Baum for helpful discussion.

## Author contributions

R.M. and N.I. designed the project. J.K., A.A.W, and R.M. performed experiments and analyzed data. J.K. and R.M. wrote the paper with comments from A.A.W. and N.I.

