## Supplementary Information for "Emergent primordial heritability in short random RNA pools"

**This file includes:**
Figs. S1 to S13

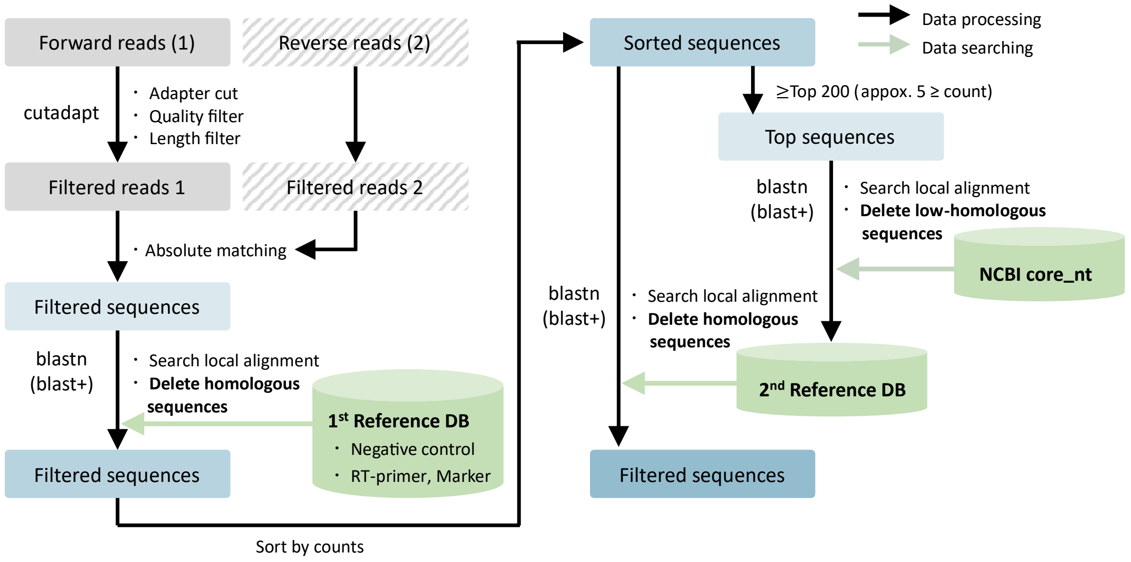


**Figure S1.** Overview of sequence processing. After quality filtering, HTS reads were further filtered through homology search against multiple custom databases (DBs).

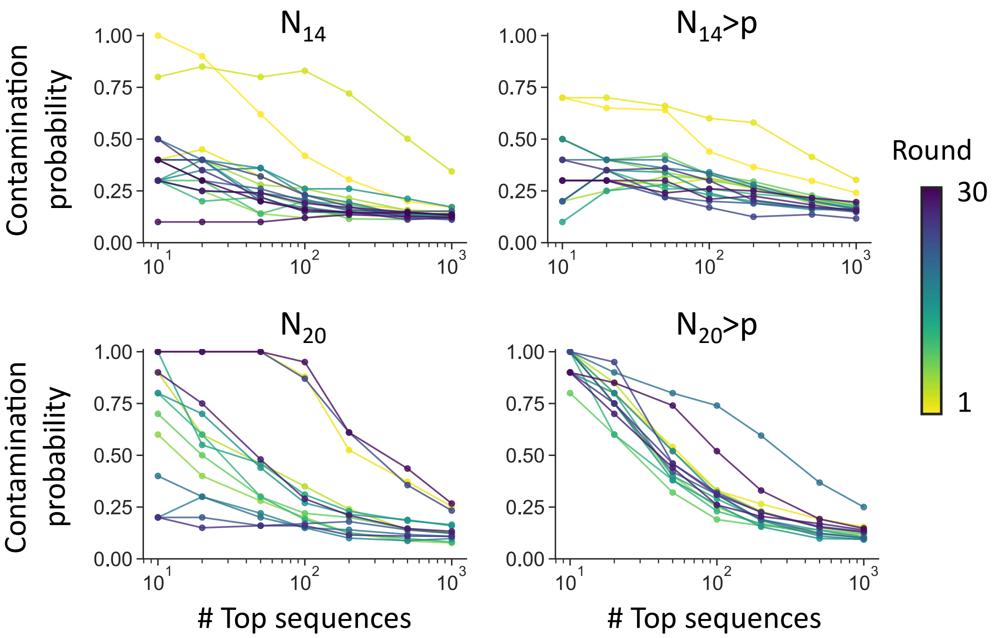


**Figure S2** Identification of potential sequence contaminants. Probability of potential contamination among the most enriched sequences in each population, based on homology searches against the NCBI core_nt database.

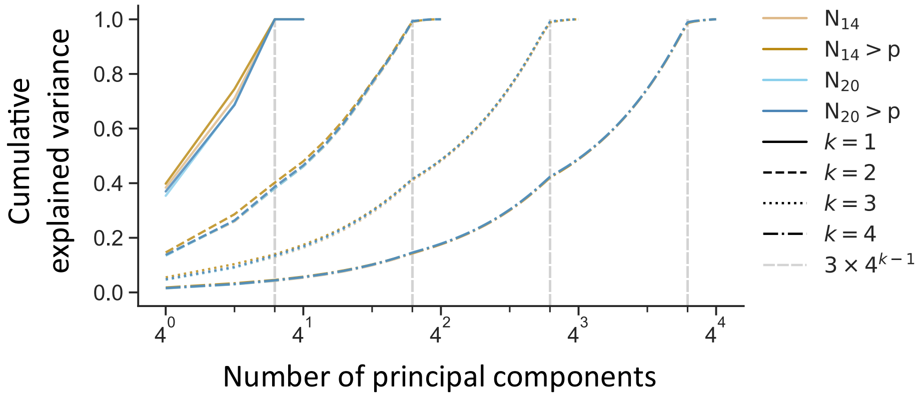


**Figure S3.** Cumulative explained variance of PCA for *k*-mer frequency vectors. Cumulative explained variance as a function of the number of principal components is shown. Values represent the mean of 10 independent PCA analyses, each performed using datasets generated by random sampling of 100,000 sequences from each round. Line colors and line styles represent analyzed pools and *k*, respectively.

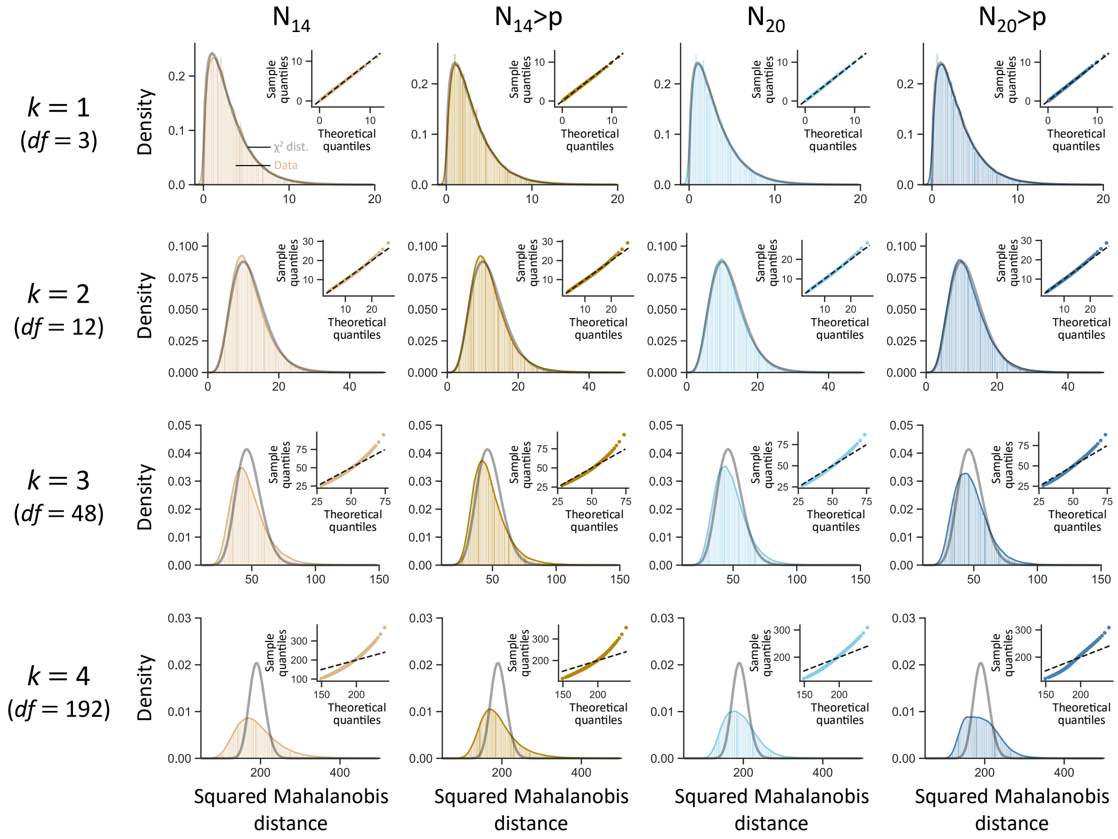


**Figure S4.** Assessment of multivariate normality of principal component scores. Squared Mahalanobis distances of the first $3 \times 4^{k\text{-}1}$ principal component scores ($k$ = 1–4) for $N_{14}$-, $N_{14}$>p-, $N_{20}$-, and $N_{20}$>p-derived products were compared with chi-squared distributions with $3 \times 4^{k\text{-}1}$ degrees of freedom (*df*) (grey lines). Data from products at the initial rounds are shown, although qualitatively similar distributions were obtained for products from other rounds. The distribution of squared Mahalanobis distances closely matched the theoretical chi-squared distribution for $k$ = 1–3. Insets show QQ plots, where agreement between empirical and theoretical quantiles indicates that the principal component scores approximately follow a multivariate normal distribution. Similar results were obtained across replicates based on at least ten independent random samplings of 100,000 sequences.

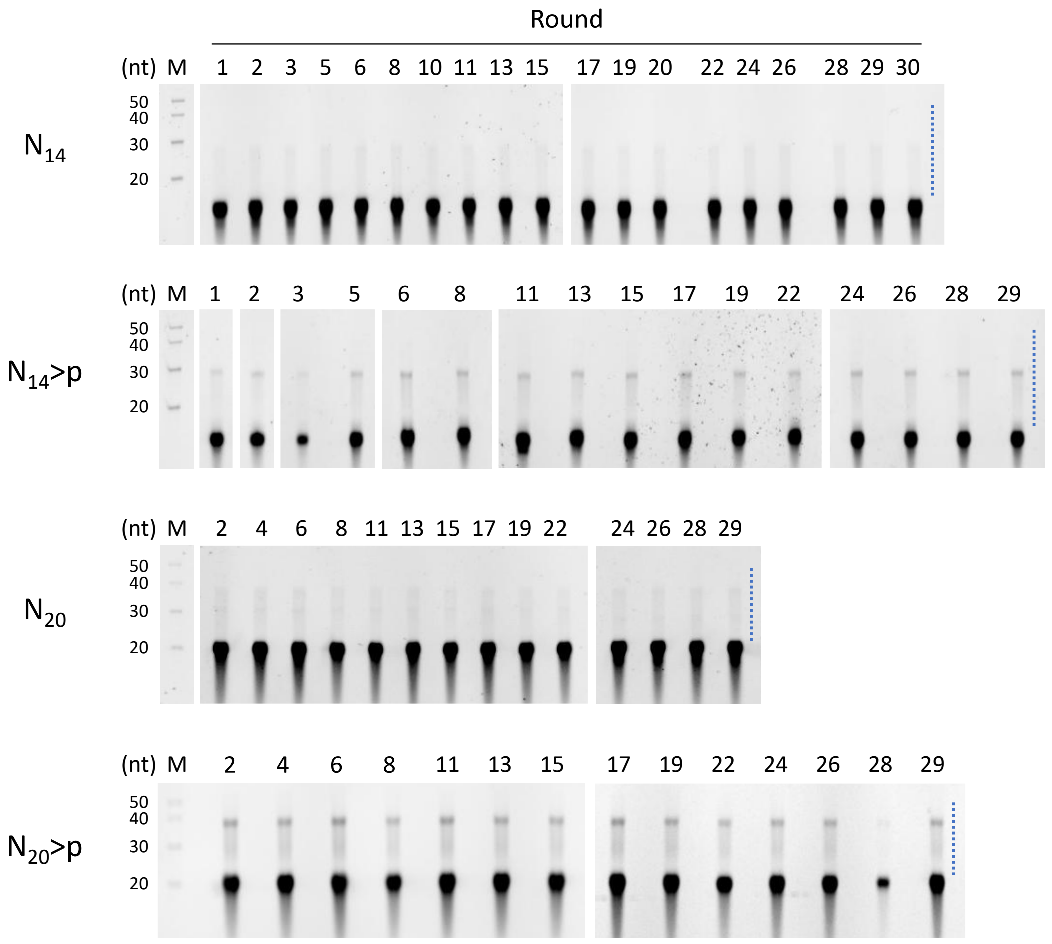


**Figure S5.** Denaturing PAGE analysis during serial transfer of short random RNA pools. RNA mixtures from $N_{14}$, $N_{14}$>p, $N_{20}$, and $N_{20}$>p pools after 2-day incubation at each round were analyzed by 20% denaturing PAGE. For $N_{20}$ and $N_{20}$>p pools, terminal phosphates were removed prior to electrophoresis. RNA products indicated by the dotted lines were excised and subjected to RT-PCR.

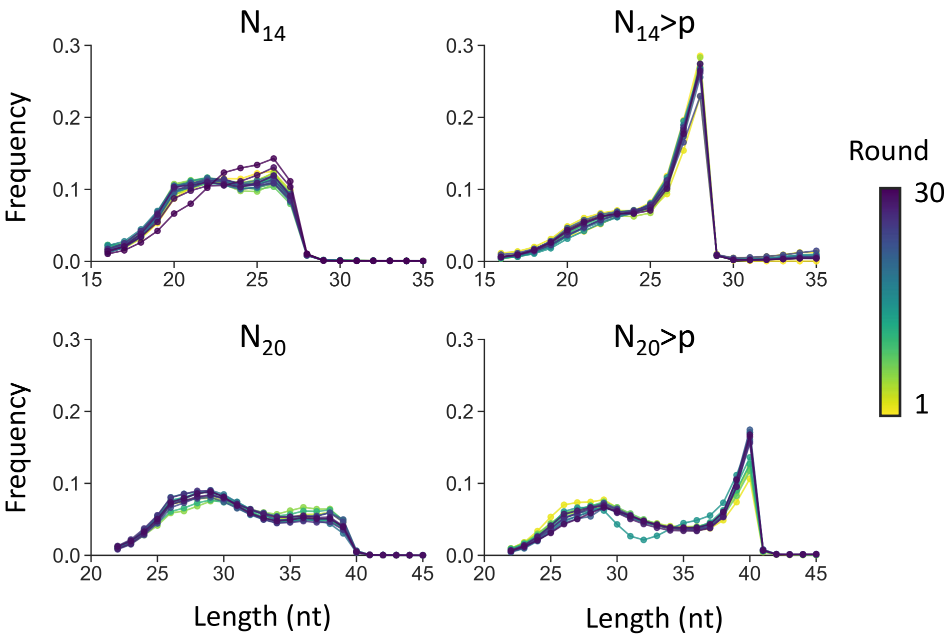


**Figure S6.** Product length distributions across all rounds. Products one nucleotide longer than the original lengths (i.e., 15- or 21-mers) were excluded as they may have arisen from the addition of an extra nucleotide to the original RNA pools during template switching.

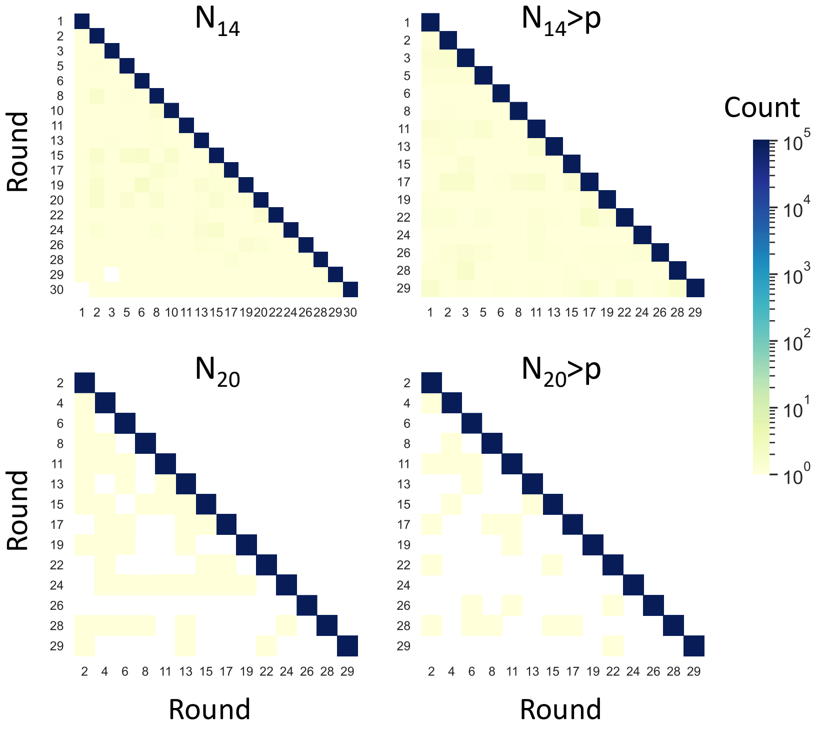


**Figure S7.** Sequence similarity among populations from different rounds. Heatmaps show the average number of identical sequences detected between any two rounds, based on random sampling of 100,000 sequences ($n$ = 10).

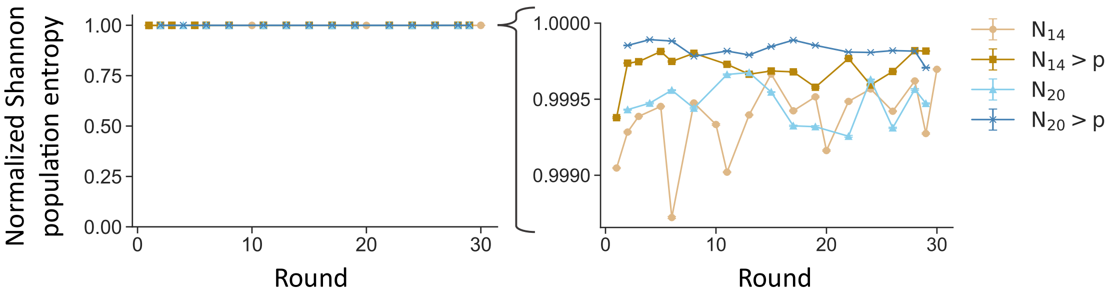


**Figure S8.** Sequence diversity of each RNA pool over rounds. Shannon population entropy was calculated as a measure of sequence diversity and normalized by the maximum entropy. Error bars indicate standard errors based on random sampling of 100,000 sequences ($n$ = 10).

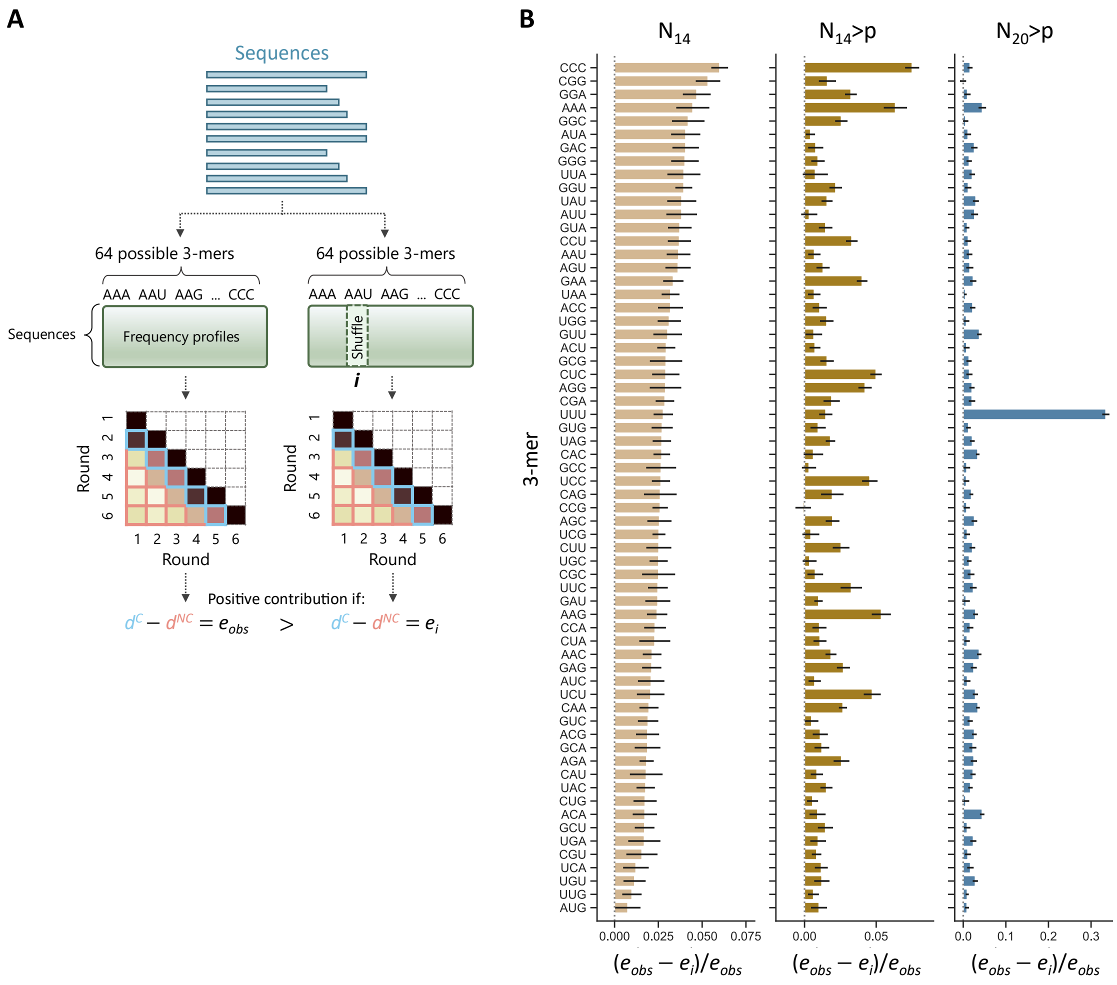


**Figure S9.** Contributions of individual 3-mers to the observed heritability. (A) Frequencies of a selected 3-mer within sequences were randomly shuffled, followed by calculation of the difference between JS divergence for consecutive rounds and that for non-consecutive rounds ($e_i$). The decrease in this difference relative to the original value ($e_{obs}$) was defined as the contribution of the corresponding 3-mer. (B) Contributions of individual 3-mers are shown for $N_{14}$-, $N_{14}$>p-, and $N_{20}$>p-derived products. Error bars indicate standard errors based on random sampling of 100,000 sequences (n = 10).

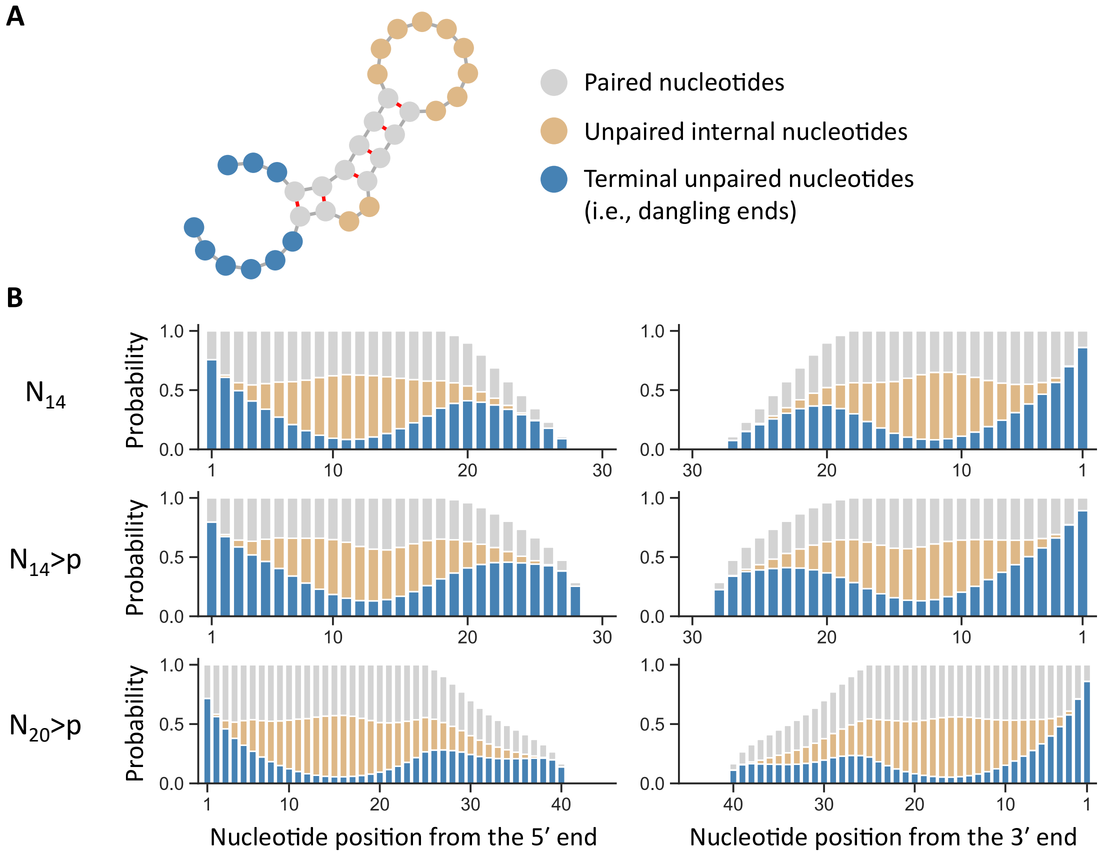


**Figure S10.** Base paring within individual sequences. (A) Individual nucleotides were categorized into three structural categories: paired nucleotides, unpaired internal nucleotides (flanked by paired nucleotides), and terminal unpaired nucleotides (i.e., dangling ends). (B) Using predicted secondary structures of products from $N_{14}$, $N_{14}$>p, and $N_{20}$>p pools, the probabilities of nucleotides assigned to each of the three categories were calculated for each position from the 5′ end (left) or 3′ end (right). For this analysis, 10,000 sequences were randomly sampled at each round and combined. Sampling was repeated 10 times, and average probabilities are shown. The same color coding as in panel A was used.

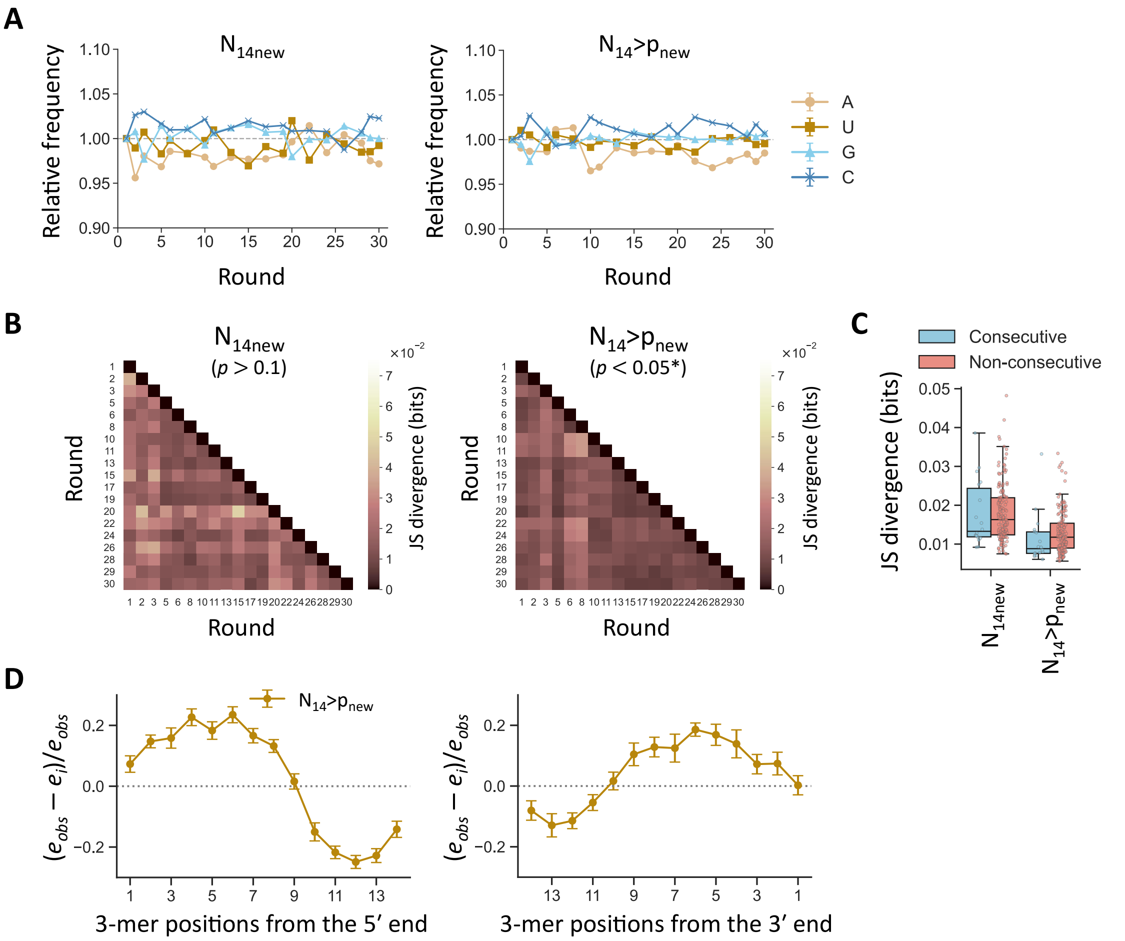


**Figure S11.** Additional transfer experiments. Two additional experiments were performed using newly prepared $N_{14}$ and $N_{14}$>p pools, denoted as $N_{14new}$ and $N_{14}$>$p_{new}$, respectively. Product sequences during the transfer experiments were analyzed as in the original experiments. (A) Changes in nucleotide composition. Error bars indicate standard errors based on random sampling of 100,000 sequences ($n$ = 10). (B) Round-by-round comparison of JS divergence based on 3-mer frequency vectors between populations at any two rounds. Average divergence from randomly sampled 100,000 sequences is shown ($n$ = 10). p-values were obtained as in Fig. 3C and are shown in parentheses; asterisks indicate $p$ < 0.05. (C) Box plots showing distributions of JS divergence between consecutive rounds and between non-consecutive rounds. Each data point corresponds to an individual divergence in panel B. (D) Position-dependent contributions of 3-mers to the observed heritable traits in $N_{14}$>$p_{new}$-derived products, quantified as changes in JS divergence after excluding specific 3-mer positions from the 5′ (left) or 3′ end (right). Error bars indicate standard errors from the same random sampling ($n$ = 10).

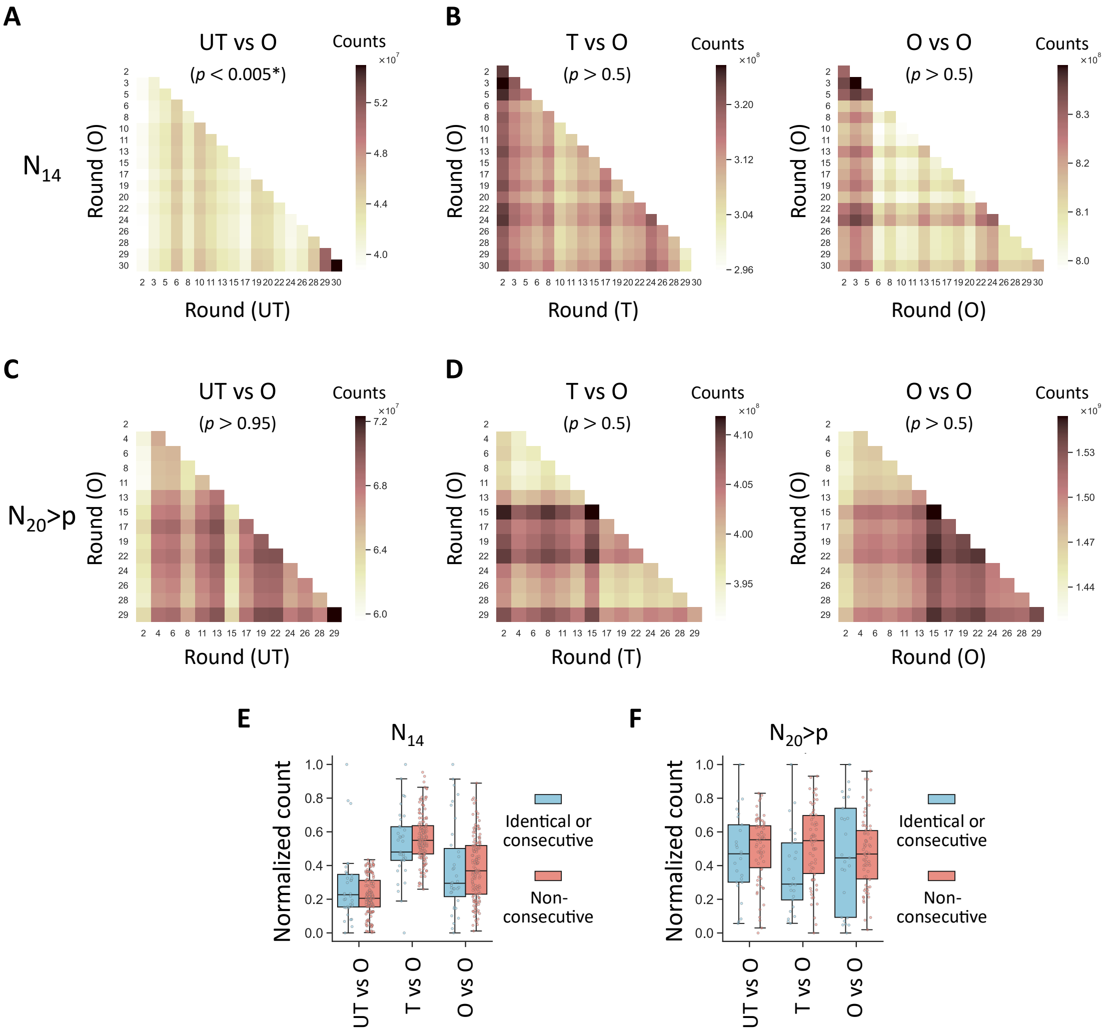


**Figure S12.** Enrichment of complementary sequences in $N_{14}$- and $N_{20}$>p-derived products. (A) Round-by-round comparison of the number of complementary 6-nt pairs between $N_{14}$-derived UT and O populations. Average counts from randomly sampled 100,000 sequences are shown ($n$ = 10). (B) Same comparison as in panel A between T and O populations (left) or between O and O populations (right). p-values calculated from the averaged matrices are shown in parentheses; asterisks indicate $p < 0.05$. (C, D) Same analyses performed for $N_{20}$>p-derived products. (E, F) Box plots showing distributions of the number of complementary 6-nt pairs after min–max scaling between identical or consecutive rounds and between non-consecutive rounds for (E) $N_{14}$- and (F) $N_{20}$>p-derived products.

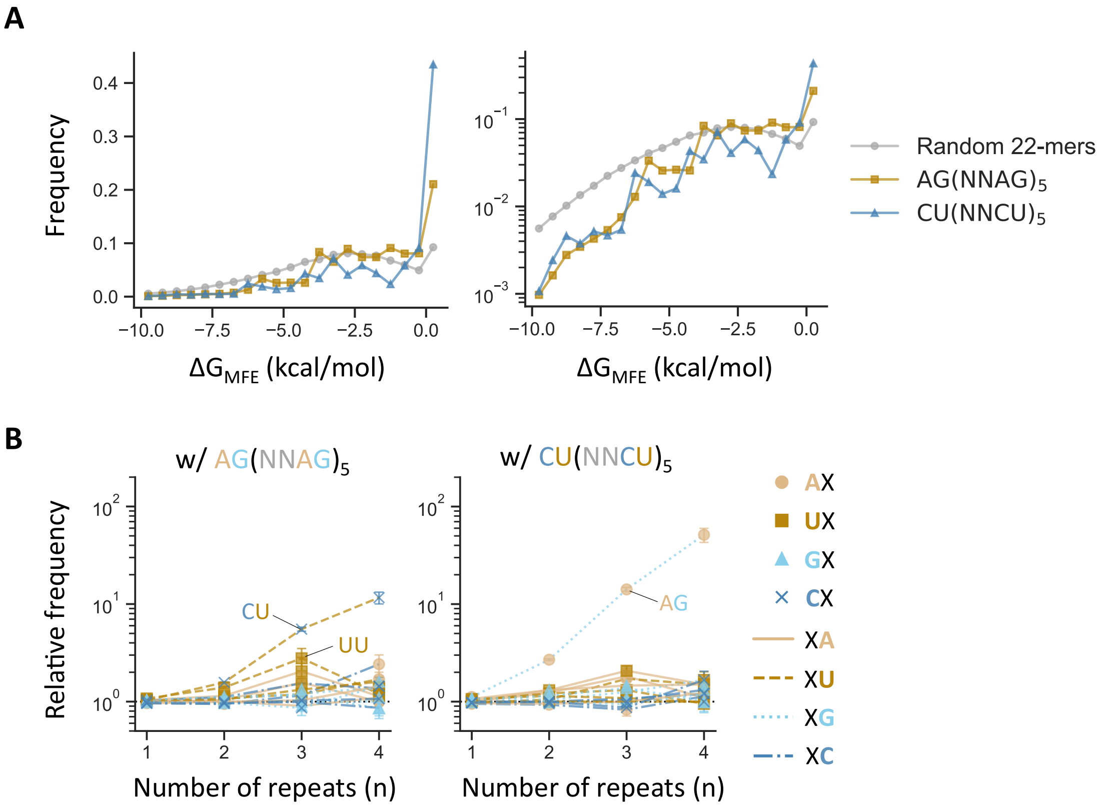


**Figure S13.** Structural propensity of semi-random RNAs and their effects at low concentrations on the dynamics of $N_{14}$>p-derived products. (A) Distribution of predicted minimum free energies ($\Delta G_{MFE}$) of fully random 22-nt RNAs and the two designed semi-random 22-nt RNAs, shown on linear (left) and log (right) scales. (B) Frequency of $XX(NNXX)_n$ motifs detected after incubating the $N_{14}$>p pool in the presence of 0.08 µM $AG(NNAG)_5$ (left) or 0.2 µM $CU(NNCU)_5$ (right), relative to that in the absence of semi-random RNAs. Different combinations of markers and line styles represent the 16 possible dimers (XX). Error bars indicate standard errors based on random sampling of 100,000 sequences (*n* = 10).
